# Molecular Strategy for Blocking Isopeptide Bond Formation in Nascent Pilin Proteins

**DOI:** 10.1101/310227

**Authors:** Jaime Andrés Rivas-Pardo, Carmen L. Badilla, Rafael Tapia-Rojo, Álvaro Alonso-Caballero, Julio M. Fernández

**Affiliations:** Department of Biological Sciences, Columbia University, New York 10027

**Keywords:** protein folding, isopeptide bond, protein mechanics, antibiotic peptide, single-molecule force spectroscopy

## Abstract

Bacteria anchor to their host cells through their adhesive pili, which must resist the large mechanical stresses induced by the host as it attempts to dislodge the pathogens. The pili of Gram-positive bacteria are constructed as a single polypeptide made of hundreds of pilin repeats, which contain intramolecular isopeptide bonds strategically located in the structure to prevent their unfolding under force, protecting the pilus from degradation by extant proteases and oxygen radicals. Here, we demonstrate the design of a short peptide that blocks the formation of the isopeptide bond present in the pilin Spy0128 from the human pathogen *Streptococcus pyogenes*, resulting in mechanically labile pilin domains. We use a combination of protein engineering and AFM force spectroscopy to demonstrate that the peptide blocks the formation of the native isopeptide bond and compromises the mechanics of the domain. While an intact Spy0128 is inextensible at any force, peptide-modified Spy0128 pilins readily unfold at very low forces, marking the abrogation of the intramolecular isopeptide bond as well as the absence of a stable pilin fold. We propose that isopeptide-blocking peptides could be further developed as a novel type of highly-specific anti-adhesive antibiotics to treat Gram-positive pathogens.

**Significance:** At the onset of an infection, Gram-positive bacteria adhere to host cells through their pili, filamentous structures built by hundreds of repeats of pilin proteins. These proteins can withstand large mechanical challenges without unfolding, remaining anchored to the host and resisting cleavage by proteases and oxygen radicals present in the targeted tissues. The key structural component that gives pilins mechanical resilience are internal isopeptide bonds, strategically placed so that pilins become inextensible structures. We target this bond by designing a blocking peptide that interferes with its formation during folding. We demonstrate that peptide-modified pilins lack mechanical stability and extend at low forces. We propose this strategy as a rational design of mechanical antibiotics, targeting the Achilles’ Heel of bacterial adhesion.

## INTRODUCTION

Bacterial infections are initiated by the attachment of bacteria to their host cells, which is mediated by specialized filamentous structures known as pili (1–3). As these adhesins are recognized virulence factors, pili have emerged as a target for the development of new vaccines and anti-adhesives (4–9). In Gram-positive bacteria, pili are assembled as a single polypeptide composed of up to hundreds of pilin protein repeats placed in tandem (10–12). Pili are known to be important factors in the virulence of Gram-positive pathogens. Indeed, failure in the proper polymerization and assembly of pili leads to diminished virulence (13–15). This central role of the pili in bacterial virulence is largely related to their unique ability to resist the mechanical challenges caused by the drag forces of mucus flow, coughing, or sneezing (16–19) (**Figure 1A**). These environmental perturbations can develop nanonewton-scale forces on a single pilus (16, 20), sufficient to unfold any protein domain stabilized through non-covalent bonds, such as networks of hydrogen bonds (21, 22). However, a key structural element of pilin proteins is the presence of intramolecular isopeptide bonds, which mechanically lock the protein structure and render it inextensible at any force (23–26). The pilus of the human pathogen *Streptococcus pyogenes* (27) is assembled as a long shaft of tandem repeats of the Spy0128 pilin protein, capped by a single adhesin Spy0125 protein. Such pili can reach several micrometers in length (10, 28, 29).

**Figure 1.**
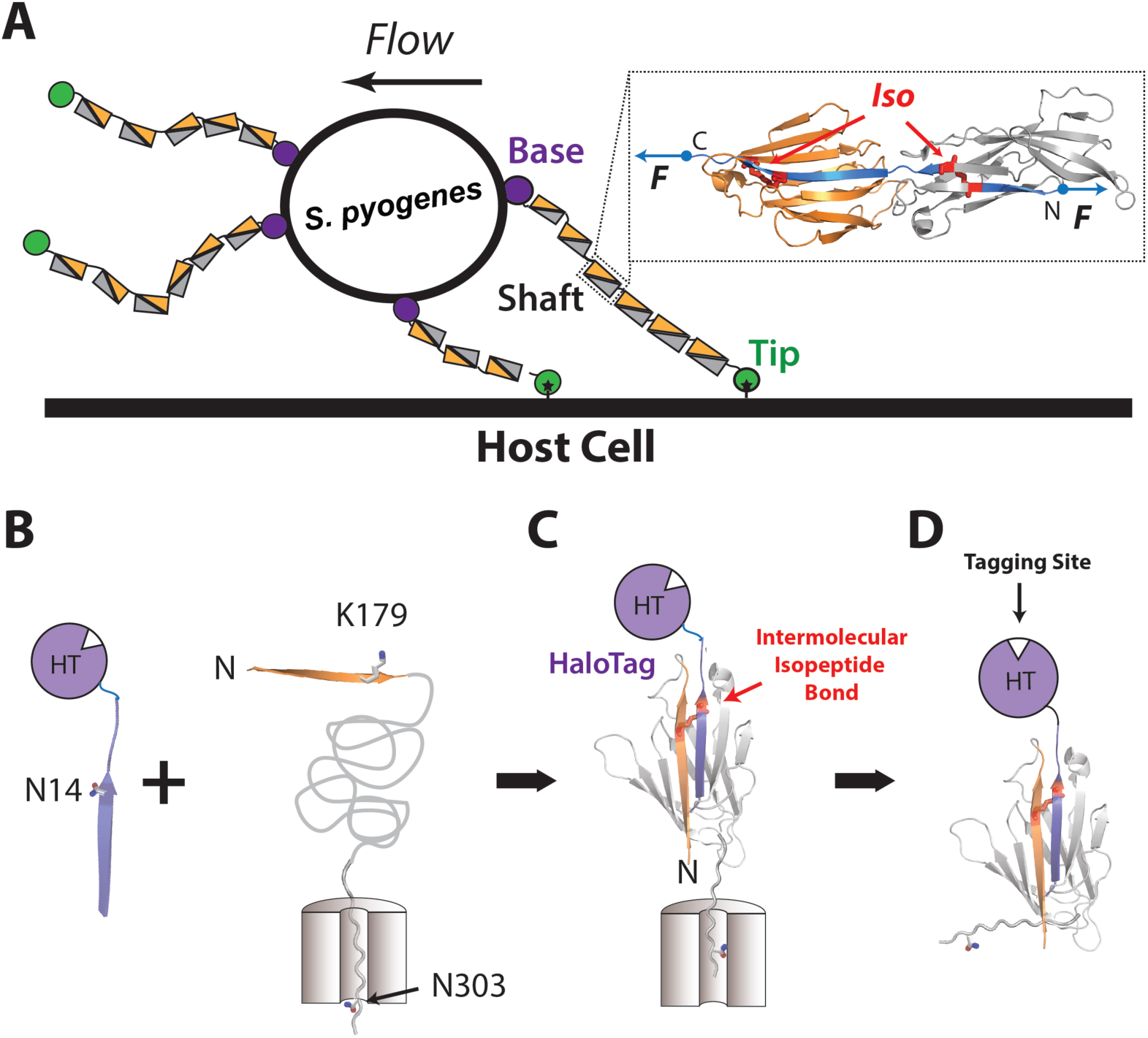
Molecular strategy for abolishing the mechanical rigidity of isopeptide stabilized pilins. **(A)** Gram positive bacteria adhere to their host cells using micrometer long pili, assembled as single modular polypeptides, which must resist shear forces in excess of 1 nN and remain immune to protease digestion. *S. pyogenes* pili are built as repeats of the pilin protein Spy0128, which is composed of two isopeptide delimited domains (inset). These isopeptide bonds are critically placed such that they shunt the force vector (blue) away from the structure, and prevent their unfolding. **(B)** In the C-terminal domain of Spy0128 (A inset; orange), the intramolecular isopeptide bond is formed between K179 and N303. We engineered a short peptide that is identical to β–strand-11, containing an asparagine residue in position 14 which matches N303 in the original structure, and competes for isopeptide formation during translation and folding. In addition, we included a HaloTag at the C terminus end of the engineered peptide. Although the channel represents the ribosomal tunnel in the experiments that we show here, it can also be understood as the bacterial translocon, through which pilin proteins are exported to the periplasm of Gram-positive bacteria, to fold and assemble to form the pilus. Being covered by a porous cellular wall, the peptide would reach the periplasmic space, and interfere with the folding of freshly exported Spy0128 proteins. **(C)** The residue N14 (peptide) competes with the emerging N303 in forming a covalent isopeptide bond, as the nascent protein folds. The result of this reaction is a domain that contains an inter-molecular isopeptide bond (K179:N14), lacking the intra-molecular isopeptide bond that gives mechanical rigidity to the pilin domain. **(D)** Modified pilin proteins can be easily detected through the use of HaloTag ligands.

The Spy0128 pilin is composed of two immunoglobulin-like domains arranged in tandem, each containing an isopeptide bond that deflects a stretching force away from the folded domain blocking their extension (**Figure 1A; inset**). For example, the C terminus Ig-like domain contains an isopeptide bond between the side chains of K179 (β–strand1) and N303 (β–strand11) (23). This intramolecular covalent bond accounts for the mechanical rigidity of the protein, as the carboxyl terminus of the C-terminal domain crosslinks to the adjacent pilin subunit at K161, which leaves the majority of the domain outside of the force pathway (23, 24) (**Figure 1A** and SI Appendix, **Fig. S1**). Hence, the isopeptide bond is responsible for the resistance of bacterial adhesion to mechanical shocks, since mechanically labile pilin proteins that extend under force are vulnerable to degradative processes (30–33). In this work, we demonstrate the design of short peptides that, by blocking the formation of the critical intramolecular isopeptide bonds, readily disrupt the mechanical properties of the Spy0128 pilin. We targeted the C-terminal domain of the Spy0128 pilin protein with a peptide—*isopeptide-blocker*—designed to mimic the β-strand11 of the pilin domain. Our strategy is simple; by presenting an exogenously added peptide that mimics β–strand11, we force emerging Spy0128 pilin proteins to form intermolecular isopeptide bonds between K179 (β–strand1) and N14 in the *isopeptide blocker* (**Figure 1B, C, D**). Timing of the intervention is critical. The intermolecular isopeptide bond between the Spy0128 pilin and the peptide must take place before the native β–strand11 of the emerging pilin can complete the fold and trigger the formation of the native isopeptide bond, after which intervention is futile. We demonstrate the crucial nature of this timing by expressing the *isopeptide-blocker* and its target Spy0128 pilins in sequence. Blockage occurs only if the isopeptide-blocker is expressed before Spy0128, resembling an antibiotic peptide which would interfere with the shaft protein before its folding and the assembly of the pilus in the periplasm. The isopeptide-blocker is Halo-tagged to label and purify modified pilin molecules, whose mechanics are thereafter investigated through atomic-force microscopy (AFM) force spectroscopy. We find that isopeptide-blocked Spy0128 pilin is extensible under force, and show reduced mechanical stability, becoming potential targets of degradation by oxidative modifications or proteases. We propose the targeting of the isopeptide bond of Gram-positive pilin molecules as a rational design of a new generation of peptide antibiotics.

## RESULTS

### Blocking intramolecular isopeptide bond formation

We designed a 19-residue long peptide, which imitates the β-strand11 of the pilin C-terminal domain (**Figure 1B**). This strand contains the asparagine residue (N303) that forms the intramolecular isopeptide bond with K179 in the N-terminal β-strand (SI Appendix, **Fig. S1**); hence, our peptide is designed to compete with the formation of the native isopeptide bond (**Figure 1B, C, D**). The isopeptide-blocker peptide is expressed as a HaloTag fusion protein, which allows for stable expression of the peptide, and for labeling and purifying modified Spy0128 proteins (**Figure 1D**) using HaloTag ligands. By using fluorescent HaloTag-ligands, we can readily identify modified Spy0128 pilin proteins in SDS-PAGE gels. Moreover, we can combine the anchoring capabilities of the HaloTag together with paramagnetic beads in order to specifically enrich the intervened fraction of Spy0128 pilin proteins recovered after their co-expression with the isopeptide-blocker peptide.

**Figure 2** shows the constructs used in our protein expression experiments: the 39 kDa Halo-tagged isopeptide-blocker (**Figure 2A**), and the 96 kDa polyprotein I91-Spy0128-I91-Spy0128-I91 (**Figure 2B**). Since the isopeptide-blocker peptide must compete with the native β-strand11 *before* the folding and formation of the native isopeptide bond, a high concentration of the isopeptide-blocker is required before the Spy0128 polyprotein folds. Thus, we co-express selectively each construct using two different inducible expression vectors within the same bacteria (*see methods*). The purified proteins are then labeled with the fluorescent Alexa488 Halo-ligand. **Figure 2C** shows an SDS-PAGE gel of the purified proteins demonstrating the labeling of the Spy0128-polyprotein by Alexa488, and its corresponding Coomassie blue on the right. Inducing the Spy0128-polyprotein alone or the polyprotein before the peptide does not result in detectable Alexa488 labeling of the Spy0128-polyprotein (**Figure 2C**, I and IA bands). However, when the peptide is expressed prior to the Spy0128, a fraction of the Spy0128-polyproteins are modified by the isopeptide-blocker resulting into two high molecular weight bands labeled with Alexa488 (**Figure 2C**, 150 kDa> AI bands <250 kDa). The Coomassie blue for total staining indicates that all three expression strategies yield comparably high levels of expression of the intact Spy0128 construct, with a molecular weight of 96 kDa (see also SI Appendix, **Fig. S2**). The HaloTag isopeptide-blocker should appear as a band at 39 kDa. However, given that we did not include a His-Tag in the isopeptide-blocker construct, the free form of this peptide is not purified together with the Spy0128-polyprotein. This was an important modification given that due to its high levels of expression, the free form of the isopeptide-blocker tends to overwhelm the Alexa488 fluorescent gels shown in **Figure 2C**. This is the reason why the 39 kDa band of the free isopeptide-blocker construct appears only faintly in these gels. The molecular weight of the fluorescent bands that appear in the gel of **Figure 2C** (i, ii, iii) and SI Appendix **Fig. S2** can be approximated by using the bands of the molecular weight standards (MW). A doubly decorated Spy0128 polyprotein should migrate at 174 kDa, while a singly decorated polyprotein at 135 kDa. Using the standards (*see methods*), we estimate that the high molecular weight bands labeled with Alexa488 correspond to 184 **(Figure 2C**; i) and 157 kDa **(Figure 2C**; ii). The branched structure of the protein constructs that result from the Halo-tagged peptides are likely to produce abnormal migration to molecular weights that do not exactly match the predicted values, which may explain these discrepancies. We also find two extra bands below and above the 75 kDa marker which do not correspond to any expected molecular weight. We speculate that they arise from the proteolytic degradation of modified Spy0128 polyproteins, which our protease inhibitors were not able to prevent during the purification. Similar proteolytic susceptibility has been demonstrated in the E258A Spy0128 protein, a mutant protein where the isopeptide of the C-terminal does not form (24, 30). In our case, this situation is even more plausible, considering that intervened Spy0128 are likely to be far more susceptible to proteolysis due to the presence of the isopeptide-blocker.

**Figure 2.**
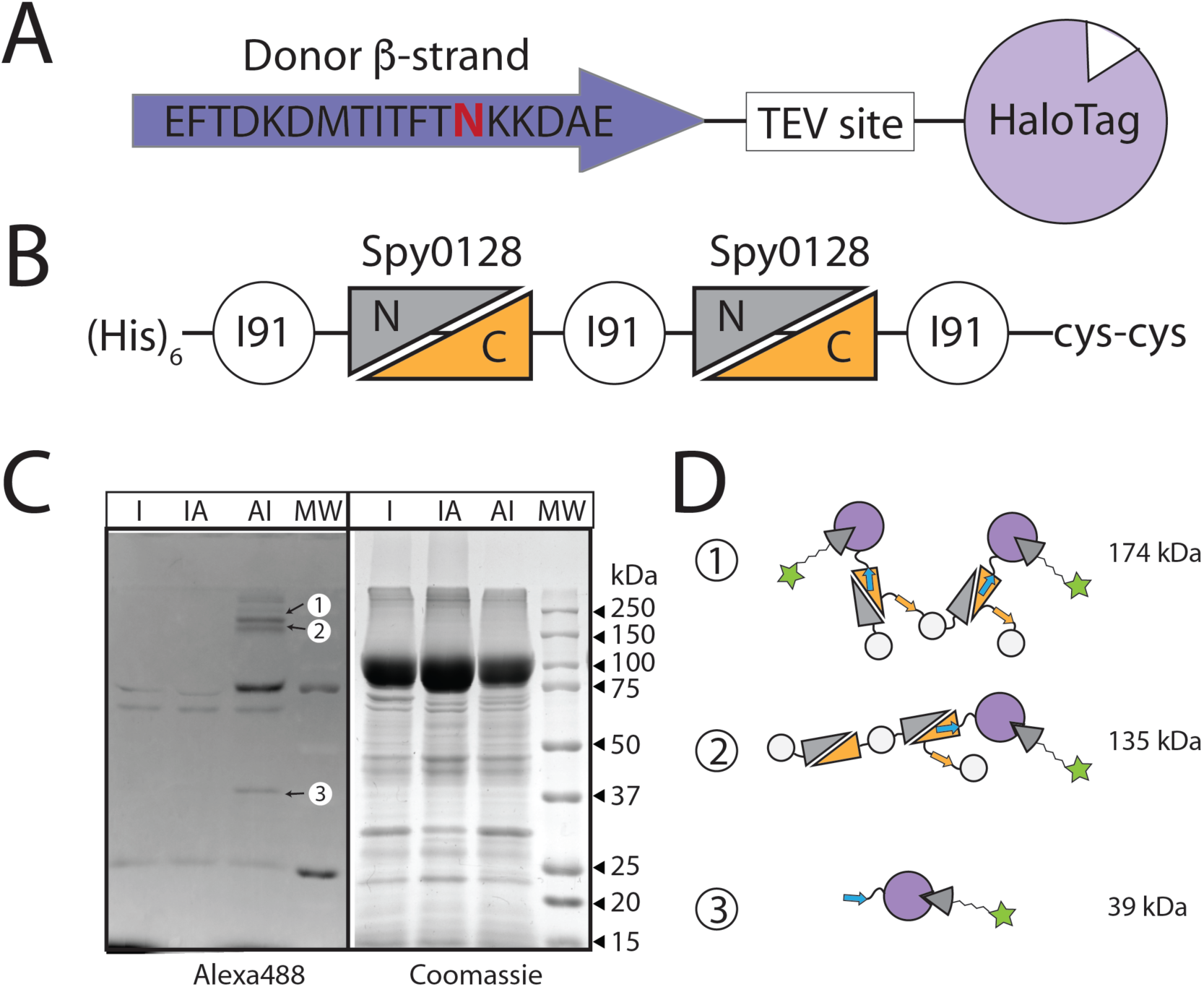
Detection of modified Spy0128 proteins using HaloTag ligands. **(A)** Design of the isopeptide-blocker. In addition to a β–strand-11 mimic (Donor β–strand), we also include a TEV site for cleaving the C terminus HaloTag used for labeling. N14 is highlighted in red. The isopeptide-blocker DNA was spliced into a vector containing an Arabinose inducer. **(B)** Design for an engineered polyprotein containing three I91 domains used as mechanical fingerprints. The I91 domains flank two Spy0128 proteins. The protein also contains two Cys residues to facilitate anchoring for the AFM experiments and a His-Tag for purification. The Spy0128 construct DNA was spliced into a vector containing an IPTG inducer. **(C)** Competent cells containing both vectors were sequentially activated and the resulting proteins were purified and labeled with Alexa488 HaloTag ligand for identification using fluorescence. Inducing with IPTG alone (I) or with IPTG followed by Arabinose (IA) does not result in high molecular fluorescent labels. By contrast, if we first induce with Arabinose (blocking peptide) followed by IPTG (Spy0128 construct), we now observe high molecular weight species labeled with Alexa488 (AI). The same gel with Coomassie staining shows similar levels of expression for all these cases. **(D)** The predicted molecular weight for a doubly modified Spy0128 construct is 174 kDa, single modified is 135 kDa, and the blocking peptide alone is 39 kDa. These bands can be found in the fluorescent gel (AI lane; i, ii, iii). Two additional fluorescent bands can be found around 75 kDa. We speculate that these bands are the result of protease degradation of modified Spy0128 constructs.

In order to demonstrate the generality of our strategy to block native isopeptide bonds, we designed two additional peptides to interfere with the structural pilin protein of a different organism, *Actynomyces oris*. In this bacterium, the shaft protein FimA shows analogous structural features with Spy0128 (34). Although FimA is a pilin protein structured as three β–sandwich domains, only the last two operate under force (34). We designed isopeptide blockers to target both the CnaA (domain N2) and the CnaB domain (domain N3) (SI Appendix, **Fig. S3**). To target the domain N2, we mimicked the β–strandI1 of the CnaA domain (Isopeptide-N), whereas for the domain N3, we mimicked the β–strandA of the CnaB domain (Isopeptide-K). We followed the same protein expression protocol used for the Spy0128 polyprotein, expressing the isopeptide-blocker before the induction of the pilin protein. SI Appendix **Fig. S3** shows that both the isopeptide-N and isopeptide-K blockers successfully decorate the FimA protein, as suggested by the bands located at 70 kDa, being 32 kDa the molecular weight of the undecorated FimA. These experiments confirm that our isopeptide-blocker strategy can be translated to shaft pilins from other Gram-positive bacteria.

### Isopeptide-blocker prevents the folding of Spy0128

We carried out AFM force spectroscopy experiments to probe the mechanics of the isopeptide-blocker modified Spy0128 protein, and resemble the putative conditions pilus proteins experience upon bacteria attachment. Specifically, we compare the unfolding mechanics of the modified pilus proteins with the intact Spy0128 and the mutant E258A, which lacks of isopeptide bond, as previously demonstrated (24). As shown in **Figure 2B**, our polyprotein constructs have two Spy0128 proteins flanked by titin I91 domains, used as fingerprint to discard spurious traces (24, 35). When pulling intact Spy0128 constructs, only the three I91 domains unfold, which are identified as unfolding events with a contour length increment of 29 nm, as determined by fits to the worm-like chain model (36), and unfolding forces of ∼200 pN (**Figure 3A**). In contrast, the mutant E258A is extensible under force, and unfolds after the I91 domains with a contour length increment of 50 nm, and unfolding forces of ∼300 pN.

**Figure 3.**
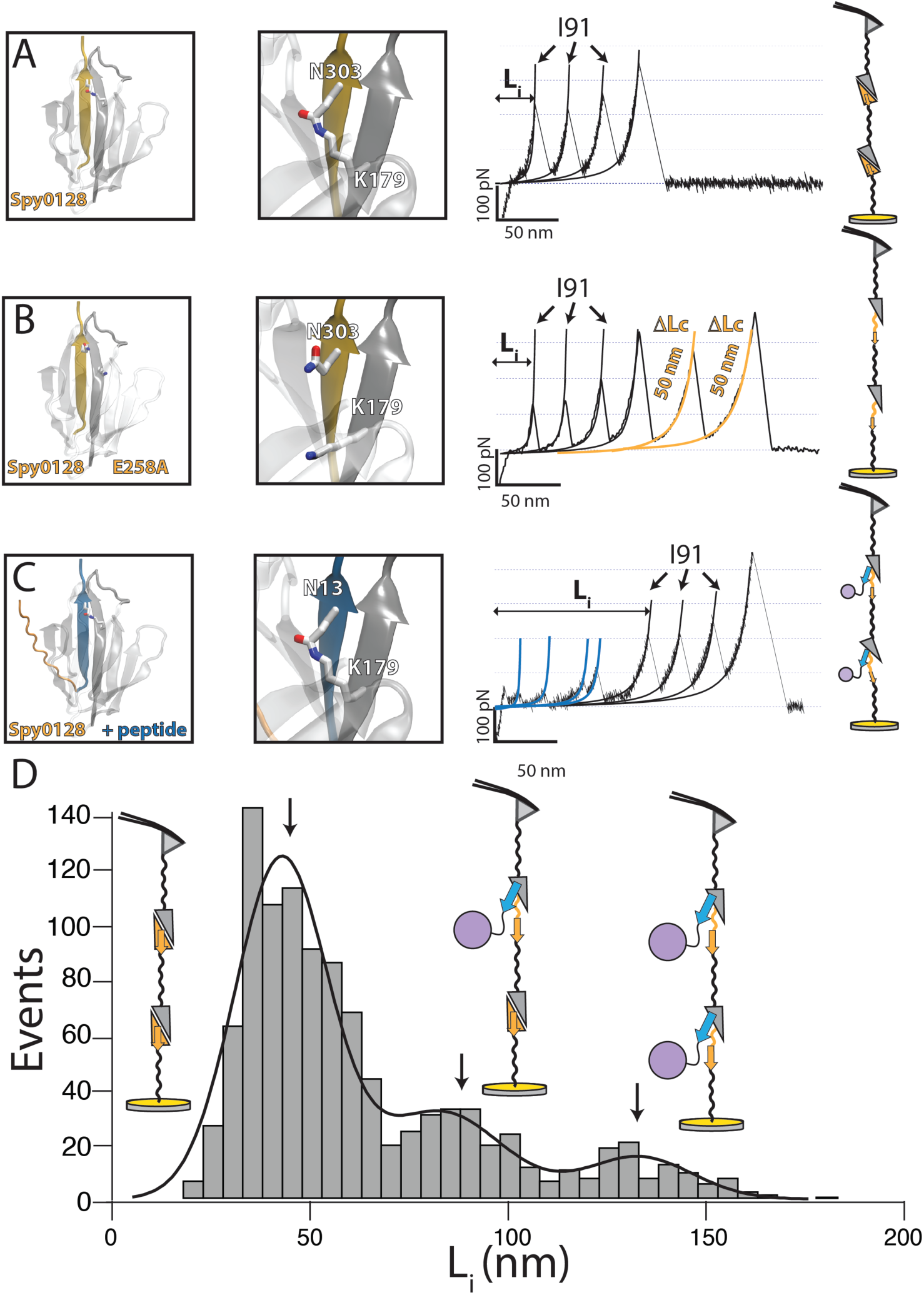
Using AFM to detect the presence, absence or the blocking of isopeptide bonds in the Spy0128 construct. **(A)** The native intra-molecular isopeptide bond in Spy0128 prevents the protein from unfolding and extending under force. Unfolding sawtooth pattern obtained by stretching the unmodified Spy0128 construct. These traces only show the three I91 fingerprints with short initial extensions (L_i_, defined as the extension to the first I91 unfolding) and contour length increments of 29 nm, measured using fits of the worm-like-chain model of polymer elasticity (solid lines). Data from (24). **(B)** Mutant Spy0128 (E258A) where the isopeptide bond is not formed. In this case, the sawtooth patterns show two additional high force unfolding events that always follow the I91 fingerprints and have large contour length increments of ∼50 nm. Data from (24). **(C)** Spy0128 constructs modified by the isopeptide-blocker show a very different type of sawtooth patterns, with long initial extensions marked by a series of random low force peaks observed always before the I91 fingerprint. **(D)** Histogram of initial extensions (L_i_) measured from sawtooth patterns obtained from Spy0128 constructs that were co-expressed with the isopeptide-blocker (n=698). A Gaussian fit (solid line) marks three distinct peaks at 41 nm, 80 nm and 130 nm. The peak at 41 nm corresponds to unmodified Spy 0128 constructs. The peaks at 80 nm and 130 nm closely correspond to the extensions predicted if one or both isopeptide bonds were blocked and the modified protein was unable to fold.

However, the isopeptide-blocker modified Spy0128 polyproteins show very different unfolding patterns, which reveal a radical change in the mechanical properties of the Spy0128. While in the former two constructs the initial extension (L_i_) never exceeds 50 nm—defined as the extension before the unfolding of the first I91 domain (35)—, modified Spy0128 proteins show initial extensions between 100 and 150 nm, which typically include several low-stability events hard to interpret systematically (**Figure 3C**). This indicates that the isopeptide-blocker, not only converts Spy0128 into an extensible protein, but also lessens its mechanical stability, when compared to the mutant E258A (**Figure 3B** and **C**).

We can take advantage of the AFM sawtooth patterns and use L_i_ as a quantitative proxy to estimate the total fraction of protein decorated with the Halo-tagged isopeptide-blocker. **Figure 3D** shows a histogram of initial extensions calculated from sawtooth patterns containing at least two I91 unfolding events (fingerprint). We distinguish three populations, centered at 41, 80, and 130 nm, respectively, which can be associated with intact Spy0128, single and double decorated, given the 50 nm contour length increment each Spy0128 renders. This allows us to estimate that 32.5% of the traces have at least one modified Spy0128, which translates into 21.1% of blocked Spy0128.

The sawtooth patterns in **Figure 3C** indicate indisputably that the mechanical stability of Spy0128 is altered, which suggests that the introduction of an extra β– strand compromises the acquisition of the folded structure of the Spy0128 C-terminal domain. Since the experiments in **Figure 3** were carried out with the purified protein, and thus the peptide contains the HaloTag, we must discard that the decreased mechanical stability is not due to steric interactions of the enzyme with the Spy0128. We carried out an extra purification step by removing the HaloTag, which also enriches the protein preparation with modified Spy0128 molecules. We incubate the polyprotein with Magne-HaloTag-beads (*see methods*), which allows us to wash out intact Spy0128 proteins, and subsequently elute the intervened polyprotein by incubating with TEV (SI Appendix, **Fig. S1**).

Interestingly, AFM experiments with the enriched fraction show equivalent unfolding patterns compared to those observed in the undigested TEV polyprotein; extended L_i_ and low mechanical stability (**Figure 3D** and **Figure 4B** and **C**). Therefore, we confirm that the diminished mechanical stability of the modified Spy0128 protein is not due to the presence of the HaloTag, but rather to the presence of the peptide. The histogram of initial extensions (**Figure 4D**) shows an increase in the population of modified Spy0128 to 60%, characterized by initial extensions of 86 and 135 nm, compatible with those described in **Figure 3**.

**Figure 4:**
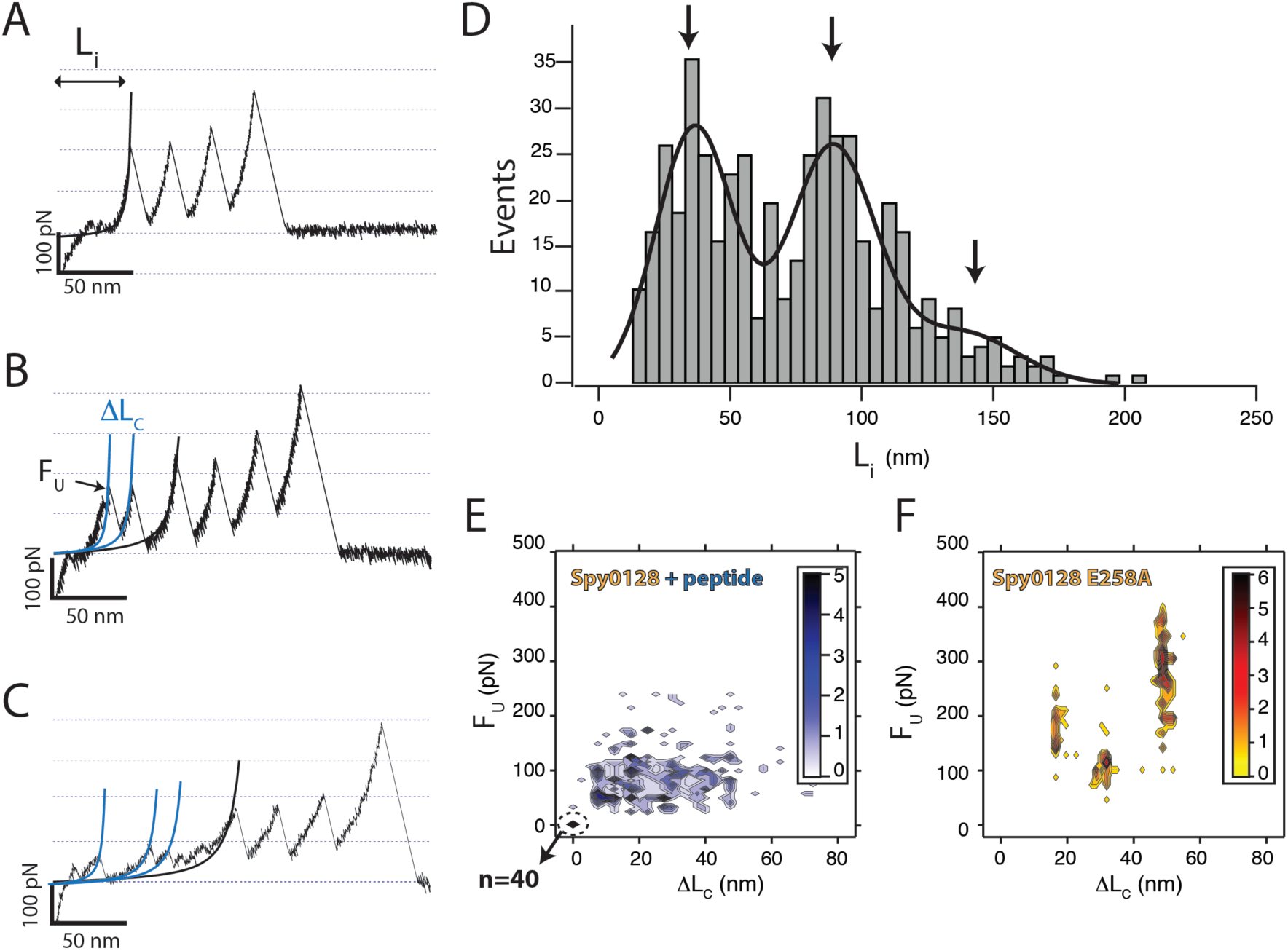
Mechanical properties of HaloTag purified Spy0128 constructs. Using Promega paramagnetic beads, we enrich the fraction of proteins that have been modified by the blocking-peptide. The proteins are subsequently detached from the beads using the TEV protease. From these purified proteins we obtain three types of sawtooth patterns: **(A)** Unmodified Spy0128 constructs with short initial extensions (L_i_ < 50 nm); **(B)** showing a single modification (L_i_ ∼50 nm - 100 nm); **(C)** or two modifications (L_i_ >100 nm). **(D)** Histogram of initial extensions (n=453). A Gaussian fit identifies three peaks at 34 nm, 86 nm, and 135 nm. **(E)** Two-dimensional histogram showing the density of unfolding forces (F_U_) versus the contour length increments (**Δ**L_c_) for the isopeptide-blocked Spy0128. Modified pilin proteins unfold at forces below 100 pN, and do not show any well-defined unfolding pathway. Remarkably, 14 % of the traces (n=40) show long initial extensions without any unfolding event, suggesting that the modified Spy0128 extends as an unstructured polymer. **(E)** Mutant Spy0128 E258A unfolds through well-defined unfolding pathways with high mechanical stability, either through a single event with F_U_∼300 pN, or through an intermediate with F_U1_∼180 pN, and F_U2_∼110 pN. Data from (24).

However, we still find a fraction of intact molecules, likely coming from contamination during the magnetic separation.

In order to characterize the mechanics of the modified Spy0128, we measured comparatively the unfolding force (F_U_) and contour length increments (ΔL_C_) of the mutant E258A, and of the unfolding events observed in the traces of the TEV-digested Spy0128 polyprotein. A contour plot of F_U_ versus ΔL_C_ for the isopeptide-blocked Spy0128 polyprotein shows scattered unfolding events with forces below 100 pN, and contour length increments ranging from 7 to 50 nm. This suggests a lack of well-defined unfolding pathways, as each modified Spy0128 shows a different unfolding pattern (**Figure 4E**). Interestingly, 14% (40 traces) of the modified Spy0128 showed large initial extensions, without any unfolding peak (ΔL_C_ = 0 nm and F_U_ = 0 pN), suggesting that the peptide can turn the Spy0128 into an unstructured polymer. In contrast, the mutant Spy0128 (**Figure 4F**) shows structured unfolding pathways: most times it unfolds through a unique step with ΔL_C_ = 50 ± 1 nm and F_U_ = 293 ± 64 pN, while in some occasions it unfolds through an intermediate ΔL_C1_ = 18 ± 1 nm and F_U1_ = 182 ± 41 pN followed by second of ΔL_C2_ = 32 ± 1 nm and F_U2_ = 111 ± 15 pN (24) (**Figure 4F**). From a structural point of view, we can hypothesize that the introduction of the isopeptide-blocker, not only abducts the native isopeptide bond, but also competes with the natural β–strand11 for the pool of non-covalent interactions that conforms the mechanical clamp, responsible for the mechanical stability of the mutant E258A. The displacement of this strand would likely give rise to steric interaction that would destabilize the fold. Remarkably, the absence of recurrent unfolding patterns in **Figure 4E** suggests that the modified Spy0128 is unable to acquire a definitive fold, but rather populates an ensemble of mechanically weak structures that results in the multi-peaked unfolding patterns.

## DISCUSSION

Gram-positive bacteria have evolved isopeptide bonds in their adhesive pili as a molecular strategy to withstand the large mechanical perturbations they experience when colonizing a host (23). These intramolecular covalent bonds clamp the shaft domains strategically to render a molecule that does not extend under nanonewton-level forces. In this environment, an extensible protein would be an immediate target for molecular degradation, such as protease digestion, or oxidative cleavage, which would compromise bacterial adhesion (31, 32). Thus, isopeptide bonds are the Achilles’ Heel of pilus mechanical integrity.

We have reported a molecular strategy to disrupt the formation of the intramolecular isopeptide bond in Spy0128, the shaft protein from *S. pyogenes*, which results in an extensible protein that unfolds under forces of few piconewton. Our strategy is inspired by the split protein technique developed by Howard and colleagues, which takes advantage of the isopeptide bond in pilin proteins to assemble molecular constructs *in vitro* (37–39). However, we target the native isopeptide bond with an engineered peptide, which must compete for its formation with the native β–strand before the folding of the pilin domain.

We suggest the design of isopeptide-blocker peptides as a novel strategy for the rational design of antibiotic peptides that target the mechanics of pilin proteins, and disrupt the early stages of bacterial infection. As we have demonstrated, the timing of our strategy is a key aspect of its success, which actually mimics the physiological process. Due to the presence of isopeptide and disulfide bonds, Gram-positive pilin proteins cannot fold in the cytoplasm, and they must translocate in the unfolded state to the periplasm (40, 41). Once in the periplasm, pilin proteins fold, form the internal covalent bonds, and polymerize with the assistance of sortases to form micron-long pili that anchor to the cell wall (11). Thus, as shown in the sequential expression of the peptide-polyprotein, nascent translocating Spy0128 would interact with the blocker peptides, forming an irreversible isopeptide bond before the protein folds; hence, displacing the native β–strand. Upon potential clinical use, mechanical antibiotics would offer an enormous specificity, given that they imitate the sequence of a β–strand of the pilin protein, and thus are exclusive for each particular pathogen. In this sense, our strategy allows combining different designs, given the specific structure of each pilin domain. While our isopeptide-blocker was the only feasible design for the Spy0128—given that the N-terminal domain does not experience force and that β–strand1 emerges the last—, FimA offered at least two targeting candidates, whose combined use would increase the fraction of extensible pilins and increase the potential effectivity of the antibiotic peptide. However, it must be noted that our inhibition strategy takes advantage of the tandem architecture of the pili—assembled from hundreds of shaft subunits—so that disrupting only a few pilin proteins would render a vulnerable anchor, whose degradation would compromise the adhesion of the entire bacteria.

Our findings provided a molecular proof of how these designed peptides target the mechanics of pilin proteins, but a potential application to clinical use would require further elaboration. Indeed, the rational design and delivery of antibiotic peptides is known to face a number of challenges, which can include proteolytic degradation, side effects, or low bioavailability (42). In this sense, the use of antimicrobial peptides is an emerging and active field, and the potential limitations are being investigated through different approaches, such as the use of nanocarriers for delivery, which has been proven to increase effectivity and reduce side effects (43, 44). Our work demonstrates that isopeptide bonds in pilin proteins are an attractive and largely unexplored target for therapeutic treatment, and future work should be aimed for *in vivo* testing of our strategy. Single cell force spectroscopy provides a natural next step to demonstrate if interfering with the mechanics of single pilin proteins scales to compromise the mechanics of single bacteria (45, 46). This would allow to propose more elaborated strategies towards a clinical implementation of isopeptide-blockers as antimicrobial peptides.

## MATERIALS AND METHODS

### Protein engineering and isopeptide-blocker design

We cloned and expressed the structural pilin protein from *Streptococcus pyogenes* (Spy0128) (23) and *Actynomyces oris* (FimA) (34). Whereas FimA was expressed as a monomer, which includes both CnaA and CnaB domains; for the Spy0128 protein, we used a previously engineered Spy0128 polyprotein (24). This polyprotein contains two Spy0128 proteins separated and flanked by three I91 titin domains, I91-Spy0128-I91-Spy0128-I91. Both constructs, FimA and Spy0128 polyprotein, were cloned in the pQE80L expression vector (Qiagen), which confers ampicillin resistance, and includes a promoter inducible by IPTG and a N-terminal His-Tag. The *isopeptide*-*blocker* was engineered by concatenating a short peptide sequence from the native β–strand region involved in the isopeptide bond formation from the Cna domain, followed by a TEV site, and the HaloTag sequence. For the Spy0128 we used the peptide –MEF<u>TDKDMTITFT</u>**<u>N</u>**<u>KKDAE</u>–, which mimics the β–strand11 of the C-terminal CnaB domain (**N**, asparagine residue that establishes the isopeptide bond). This peptide was followed by – EDIRS– (linker1), –EDLYFQS– (TEV site), –DNTTPE– (linker2), and the HaloTag. In the case of the FimA, considering that both domains experience force during the bacterial attachment(34), we targeted both the CnaA (N2 domain) and CnaB domain (N3 domain). For the N2 domain, we used –MEF<u>ARNGAIT</u>**<u>N</u>**<u>RAQVISD</u>–, which mimics the β–strandI1, penultimate β–strand of the CnaA domain, whereas for the N3 domain we used –MEF<u>WGDLLIK</u>**<u>K</u>**<u>VDNHQQG</u>–, which mimics the β–strandA, first β–strand of the CnaB domain. Both peptides were followed by – HGVRS– (linker1), TEV site, –DNTTPE– (linker2), and the HaloTag. The three isopeptide-blocker constructs were cloned in a custom modified pBAD18 expression vector (ATCC), which confers Kanamycin resistance, it is inducible by arabinose and lacks of His-Tag.

### Protein expression, purification, and labeling

The co-expression of the pilin protein and the Halo-tagged isopeptide-blocker was conducted in two stages. pQE80L and pBAD co-transformed in *E. coli* BLRDE3 pLysS cells were grown at 37° C in minimum M9CA broth (VWR) supplemented with 40 μg.mL^-1^ thymine, 2 mM magnesium sulfate, 0.1 mM calcium chloride, and 1.5 % glycerol as carbon source until the culture reached an OD_600_ of 0.5-0.6. First, the expression of the isopeptide-blocker was induced with 0.2% of arabinose for three hours at 37° C. Second, the pilin proteins were expressed in LB broth for three additional hours at 37° C with 1 mM IPTG. In the control, the protocol was inverted.

The pilin proteins were purified using the protocol described previously (23, 25). Briefly, after the *E. coli* cells were lysed by French press in phosphate buffer (50 mM Na_2_HPO_4_/NaH_2_PO_4_ pH 7.0, 300 mM NaCl), the proteins were purified by Ni-affinity chromatography (QIAGEN), followed by size exclusion chromatography (Superdex S200, GE healthcare) in Hepes buffer (10 mM Hepes buffer pH 7.2, 150 mM NaCl, 1 mM EDTA). The purified protein fractions were labeled with 1 mM of the fluorescent dye Alexa488-Halo-ligand (Promega) and analyzed by SDS-PAGE. The protein gels were imaged in a documentation gel station (G-box, Syngene) using epi illumination and a proper filter set. The molecular weight of the intact pilin proteins, isopeptide-blockers, and the decorated FimA and Spy0128 polyprotein, was calculated with Gel-Pro Analyzer 3.1 (Media Cybernetics) using the molecular weight standard as reference (precision plus protein dual color, Bio-Rad).

Selected Spy0128 polyprotein fractions were concentrated to a volume of ∼50 μL and incubated with 50 μL of pre-equilibrated Magne-HaloTag-beads (Promega) for >4 hours at 4° C in a tube rotator. The intact polyproteins were washed 3 times in Hepes buffer by centrifugation for 3 min at 1000 g at 4° C. The immobilized proteins were eluted by incubation of the beads with 100 units of ProTEV Plus (Promega) at 4° C in Hepes buffer overnight in a tube rotator. Finally, we applied a magnet to separate the proteins from the Magne-beads. TEV-treated fractions were stored at 4° C for further use.

#### Atomic Force Microscope Experiments

Polyproteins purified with or without the extra Magne-HaloTag-bead TEV step were incubated for 30 minutes in freshly nickel-chromium-gold evaporated glass cover slides. The experiments were conducted at room temperature using a custom built or a commercial AFM (Luigs and Neumann). Each of the silicon nitride cantilevers (MCLT) used in the pulling experiments was calibrated following the equipartition theorem (47), giving a spring constant of 10-20 pN. nm^-1^. The polyproteins were nonspecifically picked by pushing the cantilever at a force of 1-2 nN, and retracting at a pulling velocity of 400 nm.s^-1^. All the experiments were done in Hepes buffer at room temperature.

#### Single Molecule Data Analysis

We used the criteria established for chimeric protein constructions: only traces containing two or three I91 unfolding events were considered for the analysis (24, 35). As the I91 domains are flanking the Spy0128, the traces included within the results should contain the unfolding of the pilin domain. The initial extension (L_i_) of the polyprotein, the contour length increments of I91, and the pilin intermediates were analyzed using the worm-like chain model for polymer elasticity (36). Histograms were fitted using single and multiple Gaussian distributions implemented in Igor 7 (WaveMetrics).

## Author contributions

J.A.R.P. and J.M.F. designed the research project; J.A.R.P. and C.L.B. did the protein expressions and the protein labeling and TEV digestion experiments; J.A.R.P. and R.T-R. conducted and analyzed the single molecule experiments; J.A.R.P., R.T-R., A. A-C, and J.M.F. wrote the article.

## Acknowledgments

This work was supported by the National Institutes of Health, grants GM116122 and HL61228 (J.M.F.). R.T-R. thanks to Fundación Ramon Areces (Spain) for its financial support. We thank all the members of the Fernandez laboratory for their helpful comments to the project, and in particular to Paulina Ramirez for her invaluable help in the expression of the different protein constructs.

## Conflict of interest

The authors declare that they have no conflicts of interest with the contents of this article. The content is solely the responsibility of the authors and does not necessarily represent the official views of the National Institutes of Health.

